# μ-opioid receptor availability is associated with sex drive in human males

**DOI:** 10.1101/2021.04.20.440432

**Authors:** Lauri Nummenmaa, Patrick Jern, Tuulia Malén, Tatu Kantonen, Laura Pekkarinen, Lasse Lukkarinen, Lihua Sun, Pirjo Nuutila, Vesa Putkinen

## Abstract

The endogenous mu-opioid receptor (MOR) system modulates a multitude of social and reward-related functions, and exogenous opiates also influence sex drive in humans and animals. Sex drive shows substantial variation across humans, and it is possible that individual differences in MOR availability underlie interindividual of variation in human sex drive. Here we measured healthy male subjects’ (*n*=52) brain’s MOR availability with positron emission tomography (PET) using an agonist radioligand, [^11^C]carfentanil, that has high affinity for MORs. Sex drive was measured using self-reports of engaging in sexual behaviour (sex with partner and masturbating). Bayesian hierarchical regression analysis revealed that sex drive was positively associated with MOR availability in cortical and subcortical areas, notably in caudate nucleus, hippocampus, and cingulate cortices. These results were replicated in full-volume GLM analysis. These widespread effects are in line with high spatial autocorrelation in MOR expression in human brain. Complementary voxel-based morphometry analysis (*n*=108) provided limited evidence for association between sex drive and cortical density in the midcingulate cortex. We conclude that endogenous MOR tone is associated with individual differences in sex drive in human males.

## Introduction

Endogenous opioids modulate a range of behaviors ranging from analgesia to socioemotional processes and pleasure (Nummenmaa & Tuominen, 2018). Although dopamine is the principal neurotransmitter responsible for reward processing, murine models show that opioids produce reward independent of dopamine (Hnasko et al., 2005). In animals, μ-opioid receptor (MOR) stimulation of the nucleus accumbens increases both incentive motivation and consummatory rewards (Berridge et al., 2010; DiFeliceantonio & Berridge, 2016; Peciña & Berridge, 2013), and injection of μ-opioid agonists into the mesolimbic reward system induces reward (Bozarth & Wise, 1981). Molecular imaging studies in humans have further demonstrated central opioidergic activation following administration of various rewards ranging from feeding to social contact and exercise-induced “runner’s high” (Boecker et al., 2008; Burghardt et al., 2015; Manninen et al., 2017). Sex is one of the most potent rewards for humans, given that copulation may lead to reproduction. Human sex drive varies both between sexes as well as between and within individuals (Baumeister et al., 2001; Twenge et al., 2017), and multiple lines of evidence suggest that the MOR system could be involved in maintenance of human sex drive (Pfaus & Gorzalka, 1987).

Opioid receptors (OR) are widely expressed in the complex neurocircuitry that underlies sexual behavior (Le Merrer et al., 2009). Yet, the exact role of OR agonists and antagonists in exciting and inhibiting sexual behaviors is complex with nuanced differences across species and conditions. In a fashion similar to that of having sex, opioid agonists may increase pleasure and liking, and the euphoric sensations following opioid administration in drug addicts has sometimes been called “pharmacogenic orgasm” (Chessick, 1960). Microstimulation studies in mice have found that injecting opioids in the medial preoptic area induces consummatory sexual behaviours (Hughes et al., 1990), but striatal administration yields less consistent outcomes (see review in Le Merrer et al., 2009). In rats, copulation also induces release of endogenous opioid peptides in the medial preoptic area of hypothalamus, as indexed by MOR internalization (Balfour et al., 2004; Coolen et al., 2004). Finally, some studies have shown that also opioid antagonists may promote sexual behaviour, as administration of naltrexone shortens ejaculation latency while increasing copulation rate in rats (Rodríguez-Manzo & Fernández-Guasti, 1995).

Opioids are among the most common illicit drugs in the US (Grant et al., 2016) and clinical studies suggest that long-term opioid use has inhibitory effects on sexual behaviour at multiple levels. In humans, administration of opioid agonist heroin results in acute suppression of lutenizing hormone, and subsequently lowered plasma testosterone levels (Mirin et al., 1980). Both short- and long-term use of μ-opioid receptor agonists also decrease sexual desire and pleasure (Birke et al., 2019). One meta-analysis found that over 50% of patients on methadone maintenance treatment suffer from sexual dysfunction (SD), most commonly due to decreased desire and libido (Yee et al., 2014). Comparable rates of SDs are reported for heroin and buprenorphine maintenance, and prevalence of SDs exceeds 90% for those on naltrexone maintenance (Grover et al., 2014). Additionally, meta-analyses have confirmed that opioid use is associated with erectile dysfunction (Zhao et al., 2017). Finally, there is also some evidence on the role of long-term opioid therapy on chronic pain being associated with SD (Chou et al., 2015). This may relate to the fact that the opioid system is also activated during sexual inhibition (Argiolas & Melis, 2013), thus blunting the ability of excitatory systems to be activated (Pfaus, 2009).

### The current study

Taken together, there is ample evidence suggesting that ORs may modulate sexual behaviour in humans and nonhuman animals, but the effects between human and animal studies are not always converging. Moreover, direct in vivo evidence regarding the role of OR in human sexual motivation is lacking. Here, using a cross-sectional design, we hypothesized that human sex drive is associated with endogenous MOR availability. We used positron emission tomography (PET) with radioligand [^11^C] carfentanil that has high affinity for MOR, and measured MOR availability in 52 healthy males. Because there is evidence on the relationship between sex drive and cerebral grey matter density in certain patient populations (Bloemers et al., 2014; Schmidt et al., 2017; Takeuchi et al., 2015) but limited data on healthy subjects (Takeuchi et al., 2015), we also addressed this issue as a secondary research question. To that end, we tested whether sex drive links with regionally specific alterations in cortical density using the voxel-based morphometry (VBM) approach of T1-weighted magnetic resonance imaging scans in a partially overlapping sample of 108 males. Sex drive was determined by self-reports. We show that frequency of engaging in sexual behavior (both masturbating and partnered sex) is positively associated with MOR availability in striatum, cingulum, and hippocampus, while there was only limited evidence for sex-drive dependent alterations in cortical density.

### Materials and Methods

#### Subjects

The study protocol was approved by the Turku University Hospital Clinical Research Services Board, and the study was conducted in accordance with the declaration of Helsinki. The PET sample consisted of 52 healthy males **(Table 1)** studied with high-affinity agonist radioligand [^11^C]carfentanil (Frost et al., 1985), retrieved from the AIVO (http://aivo.utu.fi) database of *in vivo* PET images hosted at the Turku PET Centre. A subset of the data were included in our previous study on MORs and subclinical depression and anxiety (Nummenmaa et al., 2020). All subjects provided written informed consent. Brain imaging data were acquired using a GE Healthcare Discovery 690 PET/CT scanner. All PET subjects and an additional sample of 56 male subjects (a total of 108 males) were scanned with Phillips Ingenuity TF PET/MR 3-T whole-body scanner using T1-weighted sequence (TR 9.8 ms, TE 4.6 ms, f lip angle 7°, 250 mm FOV, 256 × 256 reconstruction matrix). Again, all subjects gave written informed consent and completed the questionnaires as a part of the corresponding experimental protocols. Sex drive was measured with self-reported frequency of engaging in masturbation, sexual fantasies and various sexual behaviours (kissing and caressing, oral, anal, and vaginal sex) with partner (Derogatis, 1978). Each item was rated on a nine-step scale ranging from “not at all” to “more than once per day” and averaged to yield total sex drive score. To rule out potential effects of anxiety and depression on MOR and GM density (Nummenmaa et al., 2020) all subjects also completed the Beck Depression Inventory II (BDI-II; (Beck et al., 1988) and the trait anxiety scale from the state-trait anxiety inventory (STAI-X; Spielberger et al., 1970). Power analysis on prior molecular imaging studies on personality and [^11^C]carfentanil binding (Karjalainen et al., 2016; Nummenmaa et al., 2020; Nummenmaa et al., 2015; Tuominen et al., 2012) suggested that an expected effect size of *r*=0.45, a sample size of 45 subjects would be sufficient for detecting the predicted effects at power of 0.95.

**Table 1.** Subject characteristics (means and standard deviations).

|  | PET and MRI sample<br>(n = 52) | MRI only sample<br>(n = 56) |
| --- | --- | --- |
| Age | 25.7 (0.71) | 30.1(8.66) |
| Sex drive | 4.01 (1.13) | 3.60(1.05) |
| BDI-II score | 3.73 (4.37) | 8.11(7.22) |
| STAI-X score | 33.57 (7.86) | 41.34(9.66) |

#### PET and MR image preprocessing

PET images were preprocessed using the automated PET data processing pipeline Magia (Kantonen et al., 2020; Karjalainen et al., 2020) (https://github.com/tkkarjal/magia) running on MATLAB (The MathWorks, Inc., Natick, Massachusetts, United States). Radiotracer binding was quantified using nondisplaceable binding potential (*BP*_ND_), which is the ratio of specific binding to non-displaceable binding in the tissue (Innis et al., 2007). This outcome measure is not confounded by differences in peripheral distribution or radiotracer metabolism. *BP*_ND_ is traditionally interpreted by target molecule density (*B*_max_), even though [^11^C]carfentanil is also sensitive to endogenous neurotransmitter activation (Zubieta et al., 2001). Accordingly, the *BP*_ND_ for the tracer should be interpreted as density of the receptors unoccupied by endogenous ligands (i.e., receptor availability). Binding potential was calculated by applying basis function method (Gunn et al., 1997) for each voxel using the simplified reference tissue model (Lammertsma & Hume, 1996), with occipital cortex serving as the reference region (Frost et al., 1989). The parametric images were spatially normalized to MNI-space via segmentation and normalization of T1-weighted anatomical images, and finally smoothed with an 8-mm FWHM Gaussian kernel.

To assess the link between cerebral density and sex drive, we performed a complementary voxelbased morphometry (VBM) analysis of the T1 images. VBM was done with SPM12 (https://www.fil.ion.ucl.ac.uk/spm/software/spm12/), which enables automated spatial normalization, tissue classification and radio-frequency bias correction to be combined with the segmentation step. Cut-off of spatial normalization was 25 mm and medium affine regularization 0.01 was used. Following normalization and segmentation into GM and WM, a modulation step was incorporated to take into account volume changes caused by spatial normalization and to correct for the differences in total brain size across subjects. Finally, the segmented, normalized, and modulated GM images were smoothed using 8-mm FWHM Gaussian kernel.

#### Data analysis

Regional effects were estimated using Bayesian hierarchical modeling using the R package BRMS (Bürkner, 2017), which uses the efficient Markov chain Monte Carlo sampling tools of RStan (https://mc-stan.org/users/interfaces/rstan). Atlas-based ROIs were generated in the MOR-rich regions in the brain (amygdala, hippocampus, ventral striatum, dorsal caudate, thalamus, insula, orbitofrontal cortex (OFC), anterior cingulate cortex (ACC), middle cingulate cortex (MCC), and posterior cingulate cortex (PCC) using AAL (Tzourio-Mazoyer et al., 2002) and Anatomy (Eickhoff et al., 2005) toolboxes. The ROI data were analysed with R statistical software (https://cran.r-project.org). Mean regional [^11^C]carfentanil *BP*_ND_ and GM densities from VBM were extracted for each region.

We used weakly informative priors: For intercepts, we used the default of BRMS (i.e., Student’s t-distribution with scale 3 and 10 degrees of freedom). For predictors, a Gaussian distribution with standard deviation of 1 was used to provide weak regularization. The BRMS default prior half Student’s t-distribution with 3 degrees of freedom was used for standard deviations of group-level effects; BRMS automatically selects the scale parameter to improve convergence and sampling efficiency. The BRMS default prior LKJ(1) was used for correlations of group-level random effects. The ROI-level models were estimated using five chains, each of which had 1000 warmup samples and 3000 post-warmup samples, thus totaling 15000 post-warmup samples. The sampling parameters were slightly modified to facilitate convergence (*adapt_delta* = 0.99 *max_treedepth* = 20). The sampling produced no divergent iterations and the Rhats were all 1.0, suggesting that the chains converged successfully. Before model estimation, predictors were standardized to have zero mean and unit variance, thus making the regression coefficients comparable across the predictors. Binding potentials were log-transformed because posterior predictive checking (Gabry et al., 2019; Gelman et al., 2013) indicated that log-transformation significantly improves model fit. The log-transformation essentially switches the model from additive to multiplicative; it also helps in model fitting because the assumption of linear additivity works poorly when the dependent variable is restricted to positive values (Gelman & Hill, 2006).

Complementary full volume statistical analysis was performed using SPM12. The normalized and smoothed *BP*_ND_ images and GM segments were entered into separate general linear models, where they were predicted with sex drive. Age was entered into the models as nuisance covariate because aging influences both MOR availability and sex drive (Kantonen et al., 2020; Twenge et al., 2017). Statistical threshold was set at *p* < 0.05, FDR-corrected at cluster level. In a complementary methodological approach, the data were analyzed by averaging voxelwise voxelwise *BP*_ND_ / GM density within the ROls.

### Results

Sex drive was independent of the depression and anxiety scorers as well as age (*r*s < 0.2, *p*s > 0.05); depression and anxiety scores however correlated significantly as expected (*r* = 0.62, *p* < 0.001). Mean distribution of MORs is shown in **Figure 1.** Regional Bayesian analysis revealed that sex drive was in general positively associated with MOR availability **(Figure 2).** The 95% posterior intervals did not overlap zero in middle and posterior cingulate cortices, hippocampus and dorsal caudate nucleus. The 80% posterior intervals did not overlap with zero in any of the tested regions. For VBM, there was only limited evidence for sex drive dependent differences in cortical density. All the 80% posterior intervals overlapped with zero and only in MCC was there was a weak association between sex drive dependent GM density increase.

**Figure 1.**
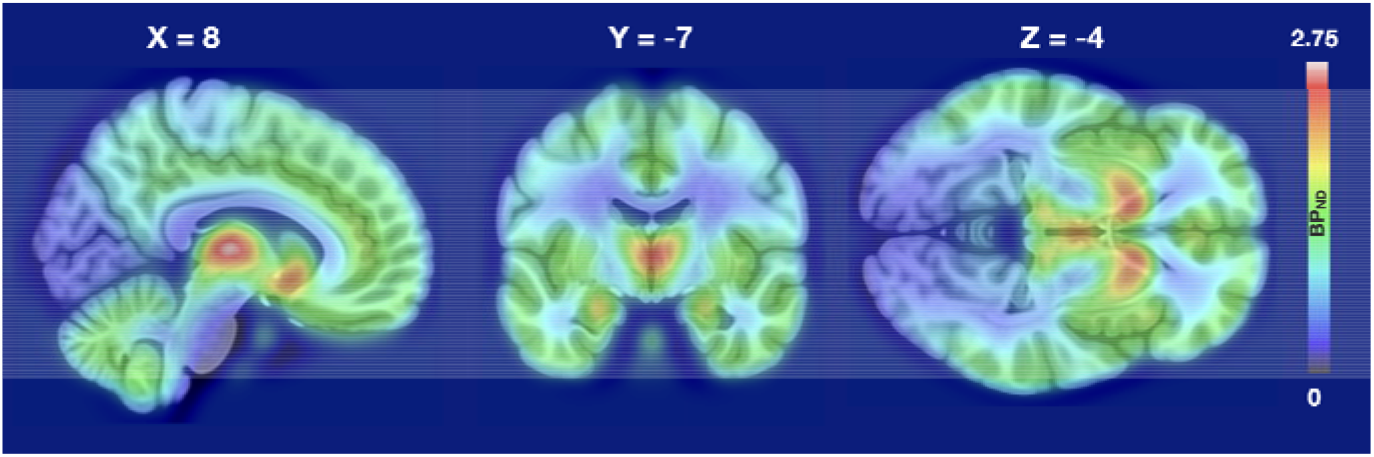
Mean distribution of [^11^C] carfentanil BP_ND_ in the sample.

**Figure 2.**
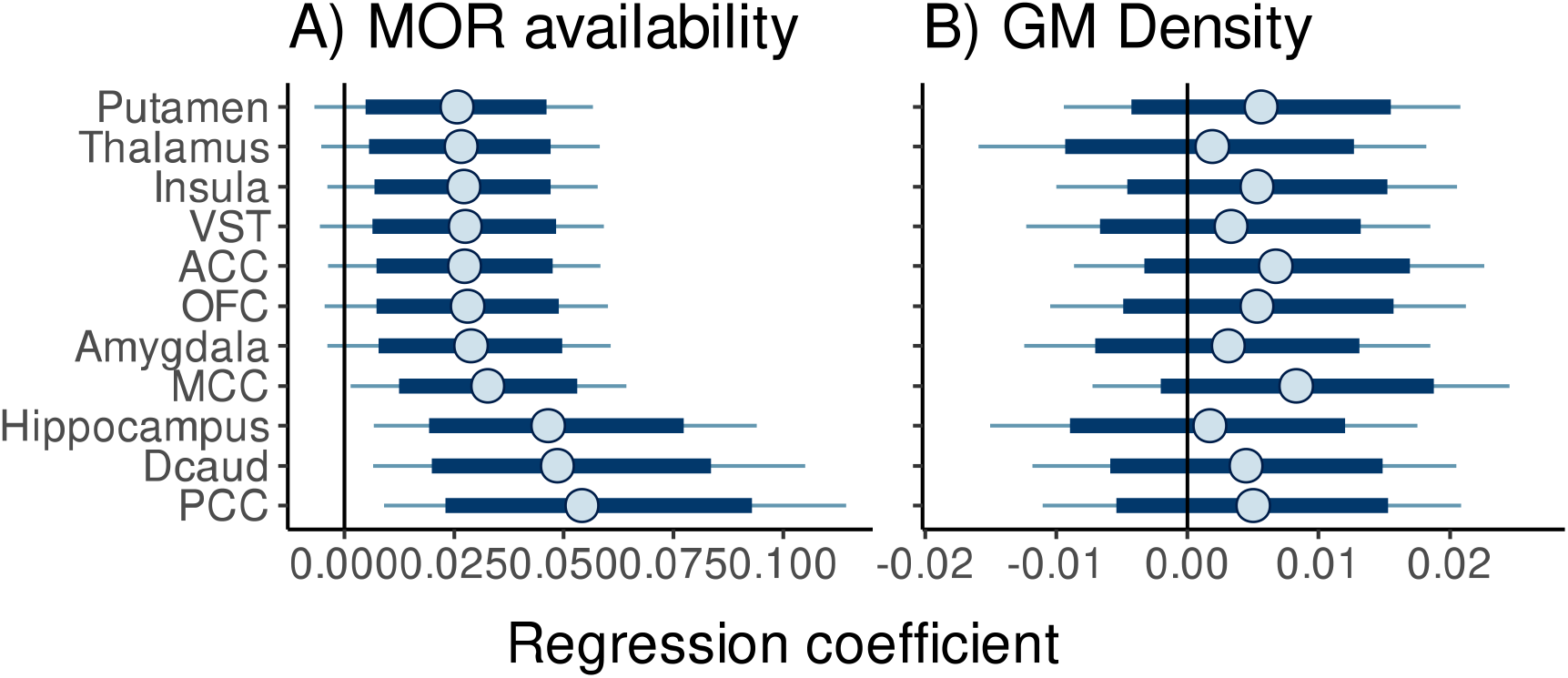
Posterior distributions of the regression coefficients for sex drive dependent variability in MOR availability (A) and cortical density (B). Thick lines show 80% and thin lines 95%posterior intervals. ACC = anterior cingulate cortex, Dcaud = Dorsal caudate nucleus, MCC = middle cingulate cortex, PFC = orbitofrontal cortex, PCC = posterior cingulate cortex, VST = ventral striatum.

The complementary full-volume SPM analysis yielded corroborating findings **(Figure 3).** Significant positive associations between sex drive and MOR availability were found in amygdala, hippocampus, cingulate cortex and ventral and dorsal striatum. Additional effects were observed in thalamus, medial and lateral frontal cortex as well as primary somatosensory and motor cortices. Again, the effects were consistently positive and when a stricter statistical threshold (*p* < .01, FDR corrected) was used, activations remained significant in the cingulate and left lateral frontal cortices.

**Figure 3.**
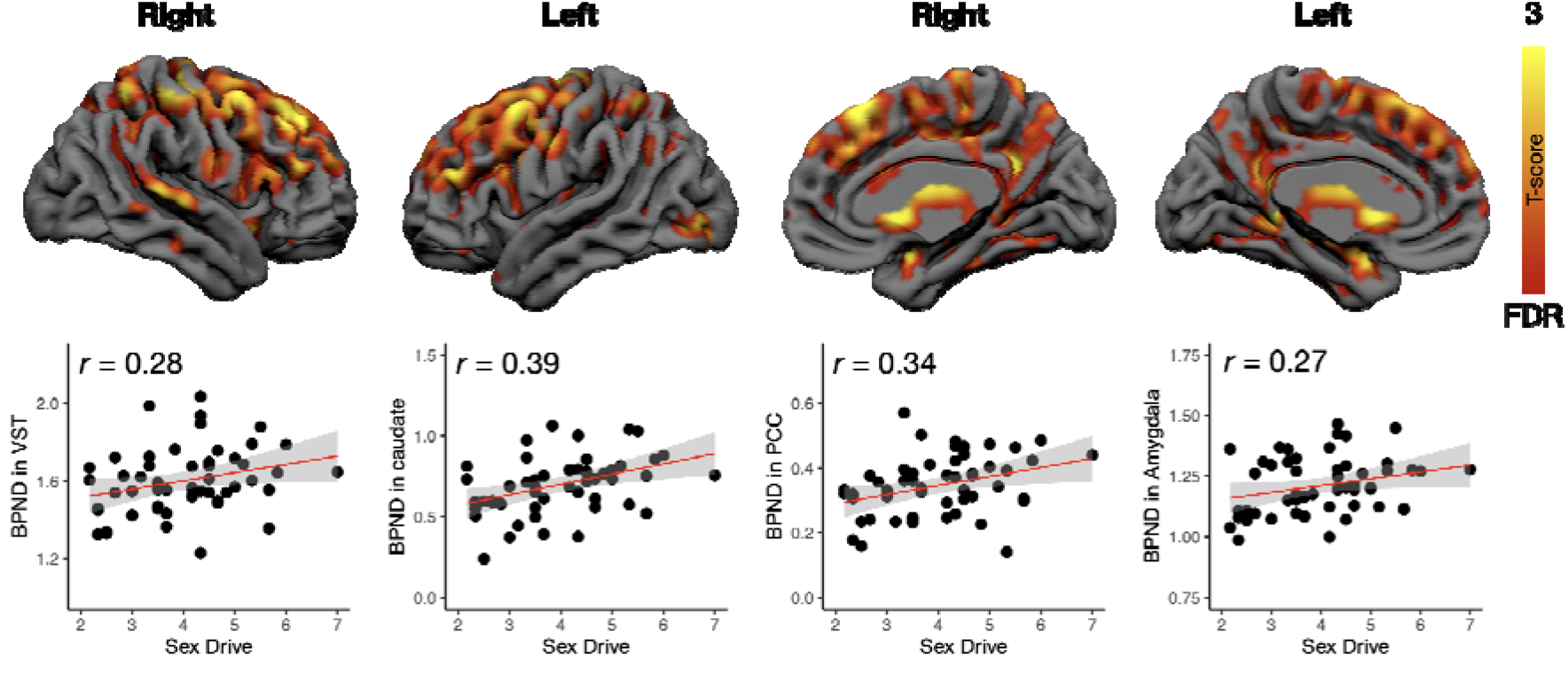
Brain regions where MOR availability was associated with sex drive. The data were thresholded at *p* < .05, FDR corrected. Scatterplots show least-squares-regression lines with 95% confidence intervals in representative regions. PCC = posterior cingulate cortex, VST = ventral striatum.

Finally, we performed full-volume GLM analysis for the GM segments. We found that sex drive was associated with increased cortical density in the anterior, middle and posterior cingulate cortex, supplementary motor cortex and primary somatosensory cortex (SI). No effects were found in extrastriatal areas **(Figure 4).** The effects in the cingulate cortex overlapped with those where sex drive dependent MOR upregulation was observed (see **Figure 3).** When stricter statistical thresholding (*p* < .01, FDR corrected) was applied, no effects remained significant.

**Figure 4.**
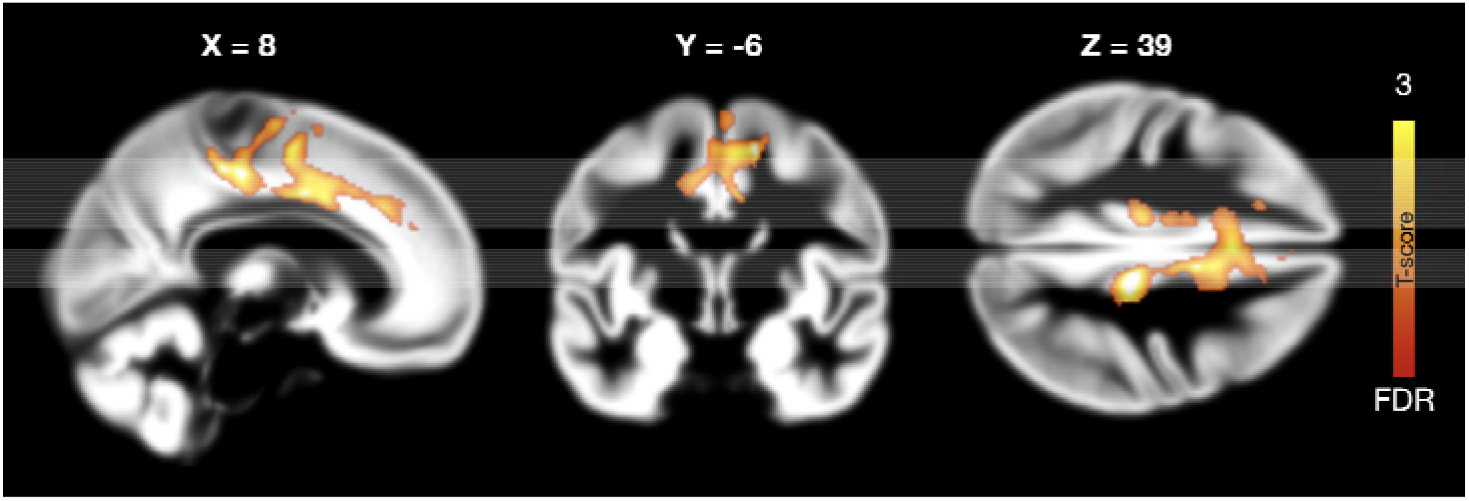
Brain regions where cortical density was associated with sex drive. The data are thresholded at p < 0.05, FDR corrected.

## Discussion

Our main finding was that male sex drive is positively associated with central opioidergic tone. The more frequently the subjects reported in engaging in sexual activities, the more μ-opioid receptors they had in the striatum, thalamus, amygdala and middle cingulate cortex. In the cingulate cortex, this effect was paralleled by increased grey matter tissue density. Our study thus demonstrates that individual differences in male sex drive are associated with availability of μ-opioid receptors, suggesting that central opioidergic mechanisms modulate not only affiliative bonding but also sexual behavior in the human male.

### Cerebral MOR availability is associated with sex drive

Sex drive had a consistent positive association with MOR availability in hippocampus, dorsal caudate and midcingulate cortices. Although the 95% posterior intervals overlapped with zero in the other tested ROI:s, the effects were systematically positive. Complementary whole-brain analysis supported sex drive dependent MOR expression in amygdala, thalamus, frontal cortex as well as primary somatosensory and motor cortices. Although the regional Bayesian and wholebrain analysis identified common regions with sex drive dependent MOR expression, the wholebrain analysis thus identified additional regions whose MOR expression was linked with sex drive. This is not unexpected, given that the whole-brain analysis approach is often more sensitive than the regional analysis, which averages data across many voxels, of which not all necessarily show similar association profiles with the predictor variables. Yet importantly, the overall pattern of results obtained with both techniques suggests a positive association between sex drive and MOR availability, with focus in the limbic and striatal regions. This general widespread effect likely reflects the high autocorrelation in MOR expression as quantified with PET (Tuominen et al., 2014).

The regions in which MOR availability was associated with sex drive are known to modulate variety of socioemotional functions (Amodio & Frith, 2006; Saarimäki et al., 2016), and they also contribute to modulating sexual behavior. While ventral and dorsal striatum modulate sexual motivation (Calabrò et al., 2019), the cingulate cortex is particularly associated with modulation of sexual drive, and meta-analyses show that anterior and middle cingulate cortices are consistently activated during sexual stimulation in humans (see review in Stoléru et al., 2012). Moreover, direct stimulation of the ACC elicits masturbation-like genital touching in the macaque (Robinson & Mishkin, 1968). Finally, the whole-brain analysis revealed sex drive dependent variability of MOR in the somatosensory cortices. Touching is a powerful way of triggering sexual arousal (Steers, 2000), and individual differences in the brevity of the sexually receptive fields of the body (“erogenous zones”) is associated with sexual drive and sexual interest (Nummenmaa et al., 2016). It is thus possible that such individual differences in the capacity for tactile sexual stimulation are dependent on MOR availability. Although hypothalamus is known to be involved in sexual functioning and that direct opioidergic stimulation of medial preoptic area induces consummatory sexual behaviour in rats (Hughes et al., 1990), we did not observe sex drive dependent effects in hypothalamic MOR availability. It is thus possible that at least in human males, hypothalamus is more involved in acute sexual motivation consummatory responses, rather than in sustained sexual drive.

To our knowledge, this is the first *in vivo* imaging study of sexual function and MOR in humans, and the present findings suggest that variation in focal MOR availability may provide an important neurochemical mechanism explaining individual differences in sex drive. Our results emphasise that this is a quantitative relationship with receptor density. It is nevertheless remarkable that MOR availability was positively rather than negatively associated with sex drive. This is a surprising observation given the general inhibitory role of OR agonist administration on sexual behaviour (see review in Le Merrer et al., 2009; Pfaus, 2009). However, comparable pattern (i.e. downregulation by agonists and positive trait correlation with MOR availability) has also been observed in the closely related phenomena of romantic and affiliative bonding, which are also modulated by MORs. Pharmacological studies in non-human primates have found that opioid antagonists promote social bonding behaviour in monkeys (Fabre-Nys et al., 1982; Graves et al., 2002; Keverne et al., 1989); conversely opioid agonists alleviate separation distress in puppies (Panksepp et al., 1978). Exogenous opioid use is also associated with lower affiliative social motivation in humans (Ross et al., 2005; Schindler et al., 2009). Paralleling the pharmacological and clinical studies, molecular imaging experiments in humans have consistently shown that MOR expression is consistently and positively associated with secure romantic and affiliative bonding (Manninen et al., 2017; Nummenmaa et al., 2015; Turtonen et al., 2021). Similarly, as sex drive linked individual differences in MOR availability, these effects are observed in the amygdala and cingulate cortices. This may reflect either opioidergic contribution to domain-general sociosexual motivation, or simply OR dependent sensitivity to rewards in general (Sander & Nummenmaa, 2021).

The more OR individuals have in the striatum, the higher pain threshold they have (Hagelberg et al., 2012). In similar vein, it is possible that individuals with high MOR availability are more tolerant to the MOR agonist driven sexual inhibition. Alternatively, it is possible that the individuals with high MOR levels simply derive more hedonic enjoyment from sexual behaviours, potentiating sex drive. Accordingly, PET imaging studies suggest that MOR availability is associated with behavioural activation system tone, which in turn is linked with appetitive motivation in general (Karjalainen et al., 2016). Both alcohol and cocaine dependence are associated with increased rather than decreased MOR availability, possibly due to reduction in endogenous opioids or upregulation of MORs (Gorelick et al., 2005; Weerts et al., 2011). It is thus possible that frequent sexual contact might similarly upregulated MOR or downregulate endogenous opioids, thus explaining the present findings.

A single baseline PET scan is not sufficient for determining the exact proportions for causal factors to the altered receptor availability which could potentially be affected by changes in receptor density, affinity or endogenous ligand binding (Henriksen & Willoch, 2008). Although [^11^C]carfentanil binding is sensitive to endogenous neurotransmitter release triggered by non-pharmacological stimulation including social contact, physical exercise, and feeding (Hiura et al., 2017; Manninen et al., 2017; Saanijoki et al., 2017; Tuulari et al., 2017) these effects are typically in the rank of 5-10% changes in the *BP*_ND_. Because [^11^C]carfentanil scans have high test-retest reproducibility (VAR□<□6%, ICC□>□0.93) (Hirvonen et al., 2009), the *BP*_ND_ from baseline [^11^C]carfentanil scans reflect predominantly tonic MOR availabilities indicating that despite transient modulations in *BP*_ND_ caused by endogenous ligands (see also Kantonen et al., 2020). In future it would be important to use the PET challenge paradigm to measure the effects of acute sexual behaviors on MOR availability.

### Sex drive and cortical density

The complementary voxel-based morphometric analysis revealed that grey matter density across the cingulate, primary somatosensory and supplementary motor cortex was negatively associated with sex drive. Although 80% posterior intervals overlapped with zero in the primary regional analysis, the overall effect of sex drive on GM density was consistently positive. The sex drive dependent effects in MOR availability and GM density overlapped in the cingulate cortex. This possibly reflects the fact that GM density estimates derived from VBM are influenced by the voxel-wise neuroreceptor densities (Manninen et al., 2021), thus the present VBM and PET data in the cingulum provide corroborative evidence on the sex drive dependent alterations in MOR expression. There is currently limited evidence on the cortical density changes associated with sexual function in healthy subjects. In one study healthy subjects’ sexual permissiveness (i.e., how acceptable people consider sexual activities in general) is negatively associated with grey matter density in amygdala in a mixed-sex sample (Takeuchi et al., 2015). Patient studies have found increased amygdala density in mixed-sex sample of subjects with compulsive sexual behavior (Schmidt et al., 2017), whereas women with hypoactive sexual desire disorder, as compared with controls, had reduced GM volume in the insula, anterior temporo-occipital and frontal cortex, as well as ACC (Bloemers et al., 2014).

### Limitations

Sex drive was based on self-reported sexual activity. These may not be perfectly accurate, as subjects may not remember their sexual activity accurately or may be reluctant to disclose their sexual behaviour. However, prior studies confirm that this kind of self-reports yield reasonably reliable results – for example partners’ retrospective reports of marital intercourse frequency are consistent (Clark & Wallin, 1964; Upchurch et al., 1991). Also, it is possible that sex drive is decoupled from the actual sexual behaviour (e.g. not engaging in sexual behaviour despite high desire to do so, or having sex without experiencing desire), yet on average the frequency of sexual behaviours is concordant with the sexual drive (Santtila et al., 2007). However, as the data were cross-sectional, we cannot conclude whether the links between MOR availability / cerebral integrity and sex drive reflect i) genetically determined individual differences in MOR availability / cortical structure (Weerts et al., 2013) contributing to increased motivation for sexual behaviour; or ii) upregulation of MOR neurotransmission and cortical density resulting from frequent sexual behaviour. Finally, our study only included young male subjects, thus the results do not necessarily generalize to older men or women due to differences in the sex-specific reproductive biology as well as sex differences in sex drive and erotic plasticity (Baumeister, 2000; Baumeister et al., 2001). Sex drive levels were in general moderately high in the sample and we did not observe associations between sex drive and age, likely due to the limited age range of the subjects. Our data cannot thus reveal whether aging and accompanying altered MOR signaling (Kantonen et al., 2020) underlies lowered sexual drive towards the old age (Lindau et al., 2007).

## Conclusions

Central opioidergic system modulates sex drive in human males. Striatal and limbic OR availability is positively associated with sex drive, and with the exception of midcingulate cortices this effect was not related to cerebral grey matter density. Although opioid system acutely suppresses sex drive (Pfaus, 2009), our study suggests that central opioidergic mechanisms modulate not only affiliative bonding but also long-term sexual behaviour in the human male.

## Authors contributions

Acquired the data: TK, VP, LS, LL, LP

Analyzed the data: LN, VP, TM

Designed the study: LN

Wrote the manuscript: all authors

## Open practices statement

None of the data or materials for the experiments reported here are available publicly because Finnish legislation does not permit redistribution of medical data such as those used in the manuscript. The experiments were not pre-registered as the study is not a part of a clinical trial.

